# Cue-related phase reset accounts for age differences in phasic alerting

**DOI:** 10.1101/413260

**Authors:** Iris Wiegand, Myriam C. Sander

## Abstract

Alertness is fundamental for the efficiency of information processing. A person’s level of alertness refers to the system’s state of general responsiveness, and can be temporarily increased by presenting a neutral warning cue shortly before an event occurs (Posner & Petersen, 1990). However, effects of alerts on subsequent stimulus processing are less consistent in older than in younger individuals. In this study, we investigated the neural underpinnings of age differences in processing of auditory alerting cues. We measured electroencephalographic power and phase locking in response to alerting cues in a visual letter report task, in which younger but not older adults showed a cue-related behavioral advantage.

Alerting cues evoked a significant increase in power as well as in inter-trial phase locking, with a maximum effect in the alpha frequency (8–12 Hz) in both age groups. Importantly, these cue-related increases in phase locking and power were stronger in older than in younger adults and were negatively correlated with the behavioral alerting effect in the older sample.

Our results are in accordance with the assumption that older adults’ neural responses may be more strongly driven by external input and less variable than younger adults’. A stronger resetting of the system in response to the auditory cue may have hindered older adults’ effective use of the warning signal to foster processing of the following visual stimulus.

## 1. Introduction

Alertness is fundamental for the efficiency of information processing as its level determines response readiness (Posner and Petersen, 1990; Sturm et al., 1999), that is, a preparatory state to react efficiently to imminent stimuli (Sturm and Willmes, 2001). A short-lived change in the cognitive system’s preparatory state can be induced by presenting a neutral warning signal (Coull et al., 2001; Posner and Petersen, 1990; Thiel and Fink, 2007), referred to as “phasic alerting effect”.

While such effects are consistently reported in younger samples, examinations of older adults are inconclusive: Some have reported preserved effects (Fernandez-Duque and Black, 2006; Haupt et al., 2018; Rabbitt, 1984), while others have shown decreased or even absent alerting effects in older age (e.g., Festa-Martino et al., 2004; Gamboz et al., 2010; Ishigami, et al., 2016; Wiegand, Petersen, Bundesen et al., 2017). The mechanisms underlying age-related decline in alerting may be elusive if age differences in processing of the alert itself are investigated in samples in which no benefit from the cue was observed (Wiegand, Petersen, Bundesen et al., 2017).

In younger adults, power modulations in the alpha band of the electroencephalogram (EEG) in response to various types of cues have been reported (Klimesch, 1999; Thut et al., 2006; Zanto et al., 2011), and were suggested to reflect a preparatory state in expectation of the stimulus (Hanslmayr et al., 2007, 2005; Jensen and Mazaheri, 2010). This alpha power modulation was shown to be reduced in older as compared to younger adults (Deiber et al., 2013). Similarly, event-related phase locking is assumed to reflect efficient temporal coordination of neural activation (Klimesch et al., 2007; Sauseng and Klimesch, 2008). Stronger phase locking indicates smaller inter-trial variance and has been associated with better performance in younger adults (Hanslmayr et al., 2005; Klimesch et al., 2004). With regard to age-related changes in phase locking, findings are inconsistent. Studies have reported reduced (Tran et al., 2016), comparable (Werkle-Bergner et al., 2012), and also increased phase locking for older relative to younger adults (Müller et al. 2009; Sander et al., 2012). Possibly, the relationship between behavioral performance and phase locking may change with age (Werkle-Bergner et al., 2012), such that high levels of inter-trial phase stability in older adults indicate a loss of complexity in the neurophysiological response and stronger entrainment by external stimulation (Garrett et al., 2013; Sander et al., 2012).

Recently, Tran and colleagues (2016) investigated age differences in alpha power and phase locking in response to a alerting cue in a visual working memory task. While they found no age differences in cue-related power, cue-related phase locking was reduced in older relative to younger adults. Furthermore, lower phase locking was associated with worse performance. However, as every trial contained a cue, the relation between cue processing and the individual alerting effect (i.e., the cue-related facilitation of stimulus processing), and potential age differences in alerting effects, could not be tested directly.

In the present study, we examined age differences in the processing of an alerting cue in a visual attention task (Wiegand, Petersen, Finke et al., 2017). Participants performed a ‘partial report task,’ in which briefly presented letter stimuli had to be discriminated (Bundesen, 1990; Duncan et al, 1999). In half of the trials, the letter display was preceded by an auditory warning cue and in the other half no cue was played. We previously demonstrated an age-specific alerting effect on visual stimulus processing in this sample (Wiegand, Petersen, Bundesen et al., 2017): Discrimination performance was significantly increased in the younger group following the warning cue, but not in the older adults. We hypothesized that the age-related decline in the alerting effect may be due to age differences in processing of the cue itself. In this study, we therefore measured power and phase locking in response to the alerting cue and examined the relations between the cue-related EEG modulations and the behavioral alerting effect in younger and older adults.

## 2. Methods

### 2.1 Participants

The participants in this study have been described earlier by Wiegand, Petersen, Finke et al. (2017). The sample consisted of 18 younger adults (YA; age: *M*_*YA*_ = 24.3 *SD*_*YA*_ = 3.1, Sex: 12 female, 8 male) and 17 older adults (OA; age: *M*_*OA*_ = 62.9 *SD*_*OA*=_7.6; Sex: 9 female, 8 male). All participants had normal or corrected-to-normal vision and none were color-blind. According to their self-report, participants were not suffering from any chronic somatic diseases, or any psychiatric or neurological impairments. The older participants were further screened for cognitive and sensory impairments. None of them exhibited symptoms of beginning dementia as indicated by scores of 26 or higher in the Mini-Mental State Examination (MMSE, Folstein et al., 1975). None of the older adults showed severe deficits in hearing [mean (*SD*) hearing thresholds (dB) were 22.7 (4.3) for 500 Hz and 23.4 (5.7) for 1000 Hz] or vision [mean (*SD*) of visual acuity 0.7 (0.2) as measured with the Snellen test]. Written informed consent according to the Declaration of Helsinki II was obtained before the experiment was carried out, and the participants received gift cards (600–700 DKK) for their participation.

### 2.2 Task

The PC-controlled experiment was conducted in a dimly lit, soundproof and electrically shielded cabin. Stimuli were presented on a CRT 17” monitor (1024×768 pixel screen resolution; 100 Hz refresh rate). Participants were seated in a comfortable chair at a viewing distance of approximately 90 cm from the screen. Each participant completed two experimental sessions on two separate days. In each of the two sessions, a total of 800 trials were run, divided into 20 blocks with 40 trials each, which lasted around 1.5 hours. Participants were given standardized written and oral instructions and example displays were presented on the screen to illustrate the task before the experiment began.

On each trial (see Fig. 1), either a single target, two targets, or a target and a distractor were presented. Two letters were presented either vertically (unilateral display) or horizontally (bilateral display), but never diagonally, resulting in 16 different display conditions. A trial began with a circle presented in the center of the screen, which participants were instructed to fixate throughout the whole trial. Then the letter array was presented on a grey background. Participants’ task was to verbally report only the red (target) letters and ignore the blue (distractor) letters. The report could be given in any (arbitrary) order and without emphasis on response speed. Participants were instructed to report only those letters they had recognized ‘fairly certainly’ and refrain from pure guessing. The experimenter entered the responses on the keyboard and pressed a button to initiate the next trial.

**Figure 1.**
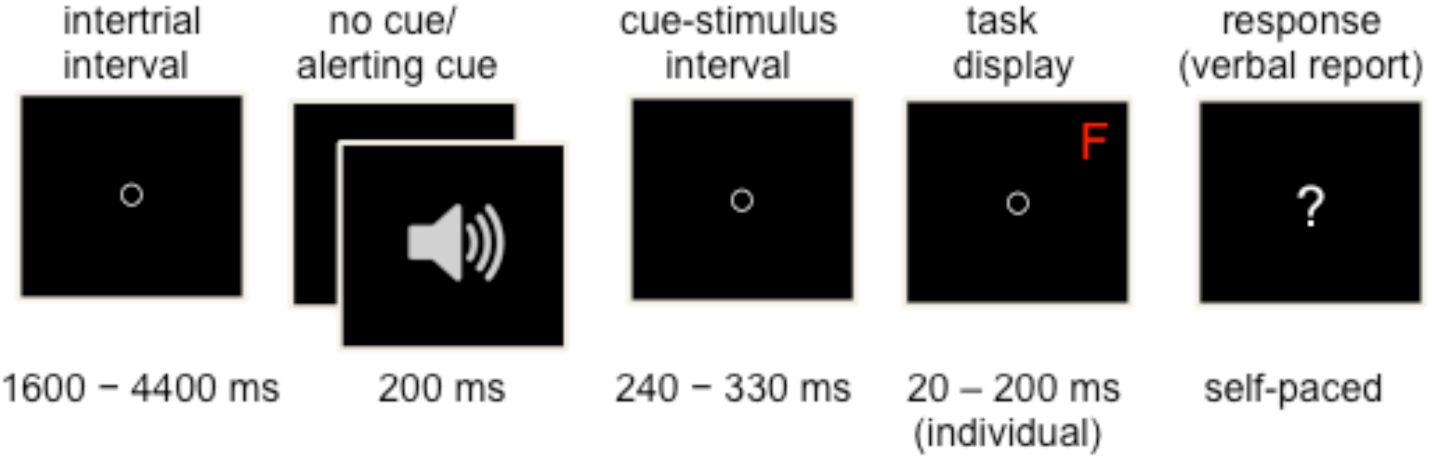
Trial procedure in the letter report task.

The exposure duration (ED) of the letter displays was determined individually in a calibration prior to the experiment (see Wiegand, Petersen, Finke et al., 2017, for details). This was done to minimize individual differences in task difficulty due to variations in perceptual threshold. EDs were chosen so that performance was 60–90% correct in single-target displays and >50% correct for individual targets within dual-target displays.

In half of the trials, randomly selected, the letter array was preceded by a loud auditory warning cue played for 200 ms. The tone was 85dB loud and varied randomly in pitch between 500 or 900 Hz to prevent habituation effects. Participants were told not to pay attention to the warning cue while performing the partial report task. The inter-trial intervals (ITIs) were drawn from a geometrical distribution with a constant hazard rate of 1/3 and a range of 1600–4400 ms using time steps of 200 ms. The cue-target intervals (CTIs) were uniformly distributed with a range of 240–330 ms using time-steps of 10 ms. In trials without a cue, time intervals identical to the CTIs were added to the ITIs to keep timing constant over conditions (see Fig. 1).

Cue and display conditions were balanced across blocks and each subject was presented with the same displays in a different random order. Letter stimuli were presented in Arial 16-point font, with equal frequencies at each of four possible display locations forming an imaginary square, with a distance of approximately 8 cm from the fixation circle. The red target color and the blue distractor color were equiluminant (2.1 cd/cm^3^, measured with ColorCAL MKII Colorimeter, Cambridge Research Systems). The letters of a given trial were randomly chosen, without replacement, from a pre-specified set (ABDEFGHJKLMNOPRSTVXZ).

### 2.3 Behavioral Data Analyses

Performance was measured as report accuracy (mean scores), that is, the rate of correctly reported individual targets in a display collapsed over upper/lower and left/right letter arrangements). We compared accuracy between trials with and without the alerting cue across all display conditions between younger and older adults via a mixed ANOVA with CUE (2) as the within- and AGE GROUP (2) as the between-subject factor.

### 2.4 EEG Recording and Preprocessing

The EEG was recorded using a Biosemi amplifier system (Amsterdam, BioSemi Active 2) from 64 active Ag-Cl electrodes mounted on an elastic cap, placed according to the International 10/10 system (American Electroencephalographic Society, 1994). Five additional electrodes were placed on the left and right mastoids, at the outer canthi of the eyes (horizontal electro-oculogramm, HEOG), and beneath the left eye (vertical electro-oculogramm, VEOG). The signal was recorded at a sampling rate of 512 Hz bandwith DC-100 Hz) and referenced online to a CMS-DRL ground, which drives the average potential (i.e., common mode voltage) as close to the AC reference voltage of the analog-to-digital box as possible (see http://biosemi.com for an explanation of the Biosemi system). The continuous signal was filtered offline with a 0.1 high-pass filter and re-referenced to the averaged mastoids and down-sampled to 128 Hz. An Infomax Independent Component Analysis (ICA; Bell and Sejnowski, 1995) using the runica algorithm implemented in EEGLAB (Delorme and Makeig, 2004) was run to identify and backtransform ocular artifacts (Jung et al., 2000). The EEG was segmented into epochs of 2 s (from −1 s prior to and 1 s following cue onset). Trials with signals exceeding +/-100 μV on any of the scalp electrodes were discarded as artifacts. The mean number of trials following artifact rejection did not differ between conditions or age groups [Younger Adults: *M*_*cue*_ = 599.89 *(SD* = 119.3), *M*_*no cue*_ 601.47 (*SD* = 120.8); Older Adults: *M*_*cue*_ *=* 591.11 (*SD* = 125.56), *M*_*no cue*_ = 589.00 (*SD* = 128.94)].

### 2.5 EEG Data Analysis

Data was analysed with Fieldtrip (Oostenveld et al. 2011), a software package developed at the F. C. Donders Centre for Cognitive Neuroimaging, Nijmegen, The Netherlands (http://fieldtrip.fcdonders.nl/) supplemented by custom-made MATLAB code (MathWorks Inc., Natick, MA, USA).

Only ICA-cleaned, artifact-free EEG trials were analyzed. Time frequency power was calculated using Hanning tapers with a fixed temporal window size of 250 ms from 4 Hz to 32 Hz in steps of 4 Hz. Following frequency decomposition, we assessed the phase stability across trials with the phase-locking index (PLI, see Lachaux et al., 1999, Tallon-Baudry et al., 1996; following Delorme and Makeig, 2004). The Fourier spectrum was divided by its amplitude, and the normalized, absolute value of the sum of angles was taken. Given that the resulting phase-locking vector varies between 0 and 1 and is potentially not normally distributed, we applied Fisher’s z-transformation. Power was log-transformed to take into account age-related changes in the 1/f ratio (Voytek et al., 2015). Power and phase locking across trials was computed for each subject, electrode, and time-frequency point, separately for cue and no-cue conditions across all trials time-locked to cue onset^1^. In a first step, we compared power and phase locking between the cue and no-cue conditions using dependent sample t-tests. Multiple comparison correction was achieved by a cluster-based permutation statistics approach (Maris and Oostenveld, 2007). Clusters were formed based on electrodes with an uncorrected p-value < 0.05 when a minimum of three adjacent electrodes reached *p*_*uncorrected*_ < 0.05. The cluster-level permutation null distribution was determined by repeating the paired t-test 1000 times, swapping the assignment for cue/no-cue conditions randomly. The summed t-values within a cluster formed the relevant test statistic. Topographical clusters were considered significant at *p* < 0.05, when the summed t-value for the true cue/no-cue assignment exceeded the 95^th^ percentile of the permutation null distribution. These analyses revealed reliable cue effects in both age groups in both neural measures. In a second step, we therefore tested directly for age differences in the amount of power increases and phase locking following the cue by comparing within-subject power and PLI differences in the cue and no-cue condition between younger and older adults. To do so, we applied independent sample t-tests corrected for multiple testing via a cluster-based permutation statistics approach with the same settings as above. Finally, using Pearson correlations, we investigated whether age differences in these neural measures were related to age differences in the behavioral cue effect.

## 3. Results

### 3.1 Effects of the Alerting Cue on Report Accuracy

The mixed ANOVA with Cue (2) as within-subject factor and Age Group (2) as between-subject factor revealed a reliable effect of the alerting Cue on accuracy *F*(1,33) = 19.1849, *p* < .001) with higher accuracy in trials with the alerting cue compared to trials without cue (*M*_*cue*_ = 0.656, *SD*_*cue*_ = 0.145; *M*_*no cue*_ = 0.635, *SD*_*no cue*_ = 0.111). Importantly, while there were no overall differences between Age Groups, *F*(1,33) = 0.60139, *p* = .44357, there was a significant interaction between Cue and Age Group, *F*(1,33) = 13.891, *p* < .001, indicating that the alerting effect had a significant effect on report accuracy in the group of younger adults (Cue: *M*_*YA*_ = 0.679, *SD*_*YA*_ = 0.10; No Cue: *M*_*YA*_ = 0.641, *SD*_*YA*_ = 0.10; *t*(17) = −7.46; *p* < .001), but not in the group of older adults (Cue: *M*_*OA*_ = 0.632, *SD*_*OA*_ = 0.13; No Cue: *M*_*OA*_ = 0.629, *SD*_*OA*_ = 0.12; *t*(16) = −0.3825; *p* = 0.71; see Fig. 2).

**Figure 2.**
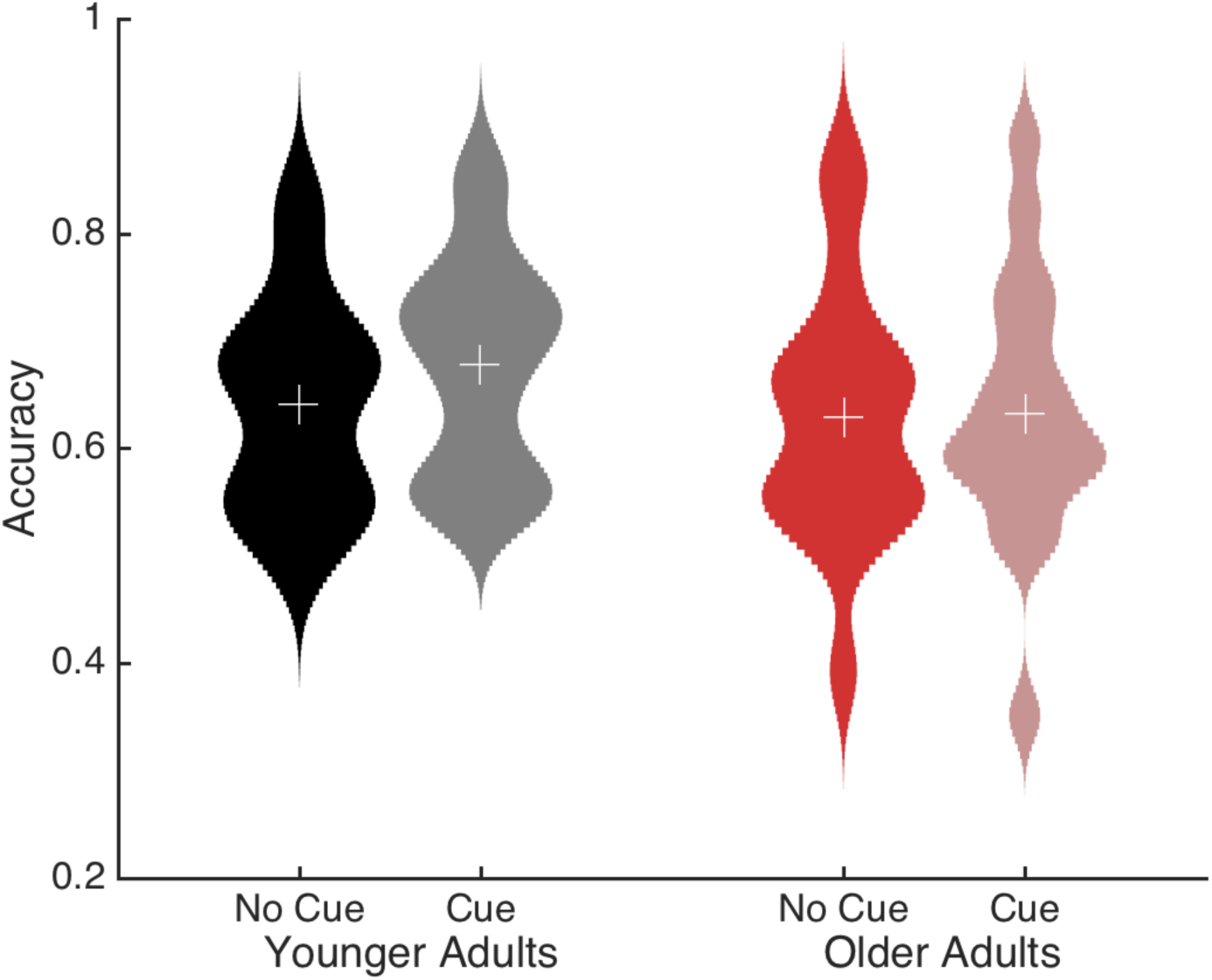
Accuracy in the letter report task for younger and older adults with and without a preceding alerting cue. Means are represented by a white cross. Whereas younger adults (YA) show better performance in the cue condition, the group of older adults (OA) does not benefit from the presentation of a cue.

### 3.2 Effects of the Alerting Cue on Ongoing Power

Cluster-based permutation statistics yielded one significant cluster of reliable power differences between cue and no-cue trials both for the younger and the older adults (*p* < 0.025). The power effects spanned frequencies in the theta, alpha, and beta range (4–20 Hz) and encompassed all electrodes. In both age groups, maximum differences between the two conditions showed a fronto-central distribution and were found in the alpha band approximately 50–200 ms after the cue onset (Fig. 3). Within the respective cluster, (log) power was higher in trials in which a cue was presented than in trials without a cue, both in younger (No Cue: *M*_*YA*_= 0.7261, *SD*_*YA*_= 0.2347; Cue: *M*_YA_= 0.8379, *SD*_YA_= 0.2314) and older adults (No Cue: *M*_OA_= 0.7284, *SD*_OA_= 0.2000; Cue: *M*_OA_= 0.8396, *SD*_OA_= 0.1789). Thus, both age groups demonstrated a reliable increase in power following the presentation of an alerting cue.

**Figure 3.**
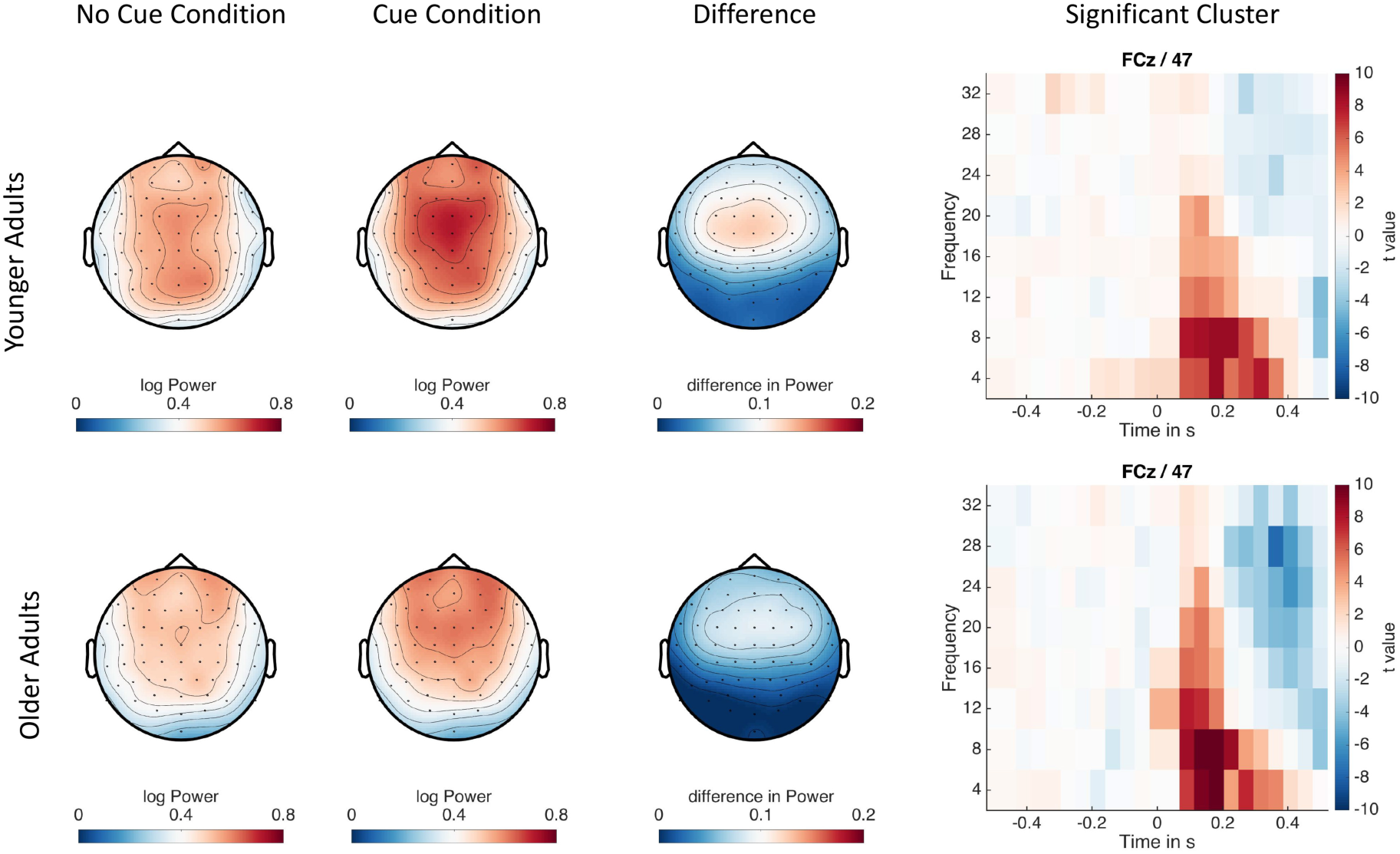
Cue-related (log-transformed) power modulation in younger (upper panel) and older adults (lower panel). Both age groups show a reliable increase in power over centro-frontal electrodes shortly after the presentation of the cue. For illustration, significant clusters at FCz are highlighted.

### 3.3 Effects of the Alerting Cue on Ongoing Phase Locking

Cluster-based permutation statistics also yielded one significant cluster of reliable PLI differences between cue and no-cue trials for both age groups (*p* < 0.025). The PLI effects were broadly distributed across all frequencies and electrodes in the post-cue time window (0–500ms), with a maximal difference over fronto-central electrodes in the alpha band around 0–300 ms after the cue onset (Fig. 4). Phase locking was stronger in trials with a cue was than in those without in both younger (No Cue: *M*_YA_= 0.0504, *SD*_YA_= 0.0075; Cue: *M*_YA_= 0.1694, *SD*_YA_= 0.0385) and older adults (No Cue: *M*_OA_= 0.0652, *SD*_OA_= 0.0171; Cue: *M*_OA_= 0.2089, *SD*_OA_= 0.0409) in the respective cluster. Thus, both age groups demonstrated reliable modulations of the neural signal following an alerting cue.

**Figure 4.**
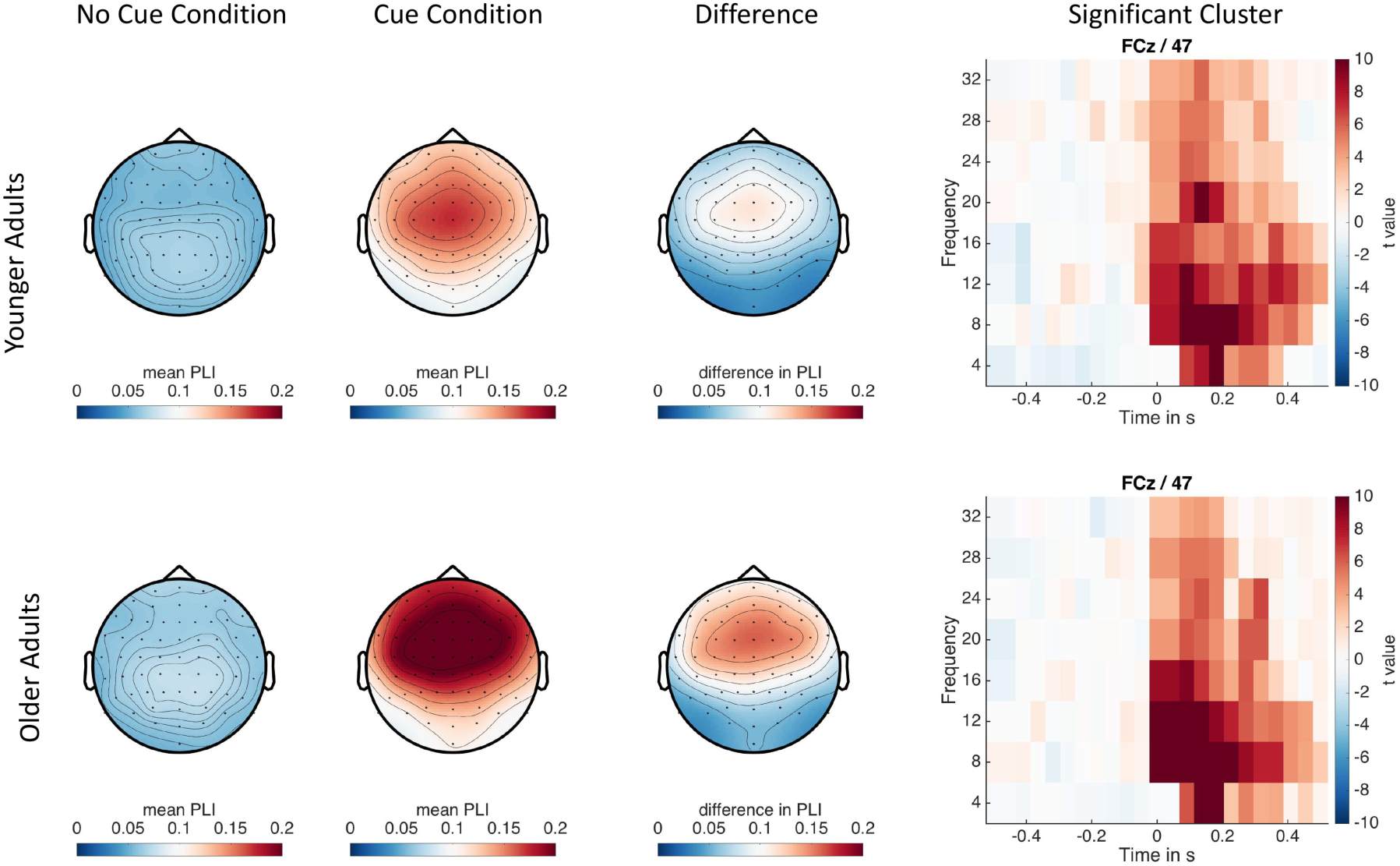
Cue-related PLI modulations in younger (upper panel) and older adults (lower panel). Both age groups display a reliable increase in phase locking over fronto-central electrodes shortly after the cue presentation. For illustration, significant clusters at FCz are highlighted.

### 3.4 Age Differences in the Effects of the Alerting Cue

Given that we observed reliable increases in both power and phase locking in both age groups following the presentation of a cue while only younger adults seemed to benefit on the behavioral level, we set out to test directly for age differences in these seemingly similar effects. We therefore subtracted power, respectively PLI in no-cue trials from power, respectively PLI, in cue trials for each participant and searched for age differences in the cue effect via cluster permutation statistics. With regard to power, this direct age comparison yielded a marginally significant cluster (*p* = 0.025) at around 50–100ms in the alpha band (8–12 Hz). Older adults showed a larger difference in power (i.e., a stronger effect of the cue) than younger adults (*M*_YA_ = 0.0978, *SD*_YA_ = 0.0554; *M*_OA_ = 0.2117, *SD*_OA_ = 0.0965).

With regard to the cue effect on the PLI, the direct age comparison revealed a clear difference (*p* < 0.025) in the alpha/beta range (8–16 Hz) in a short time window following cue onset (0–100 ms) with a fronto-central distribution (Fig. 5). Importantly, this difference was again driven by older adults showing a larger difference in phase locking (i.e., a stronger effect of the cue) than younger adults (*M*_YA_ = 0.2471, *SD*_YA_ = 0.0893; *M*_OA_ = 0.4134, *SD*_OA_ = 0.1167).

**Figure 5:**
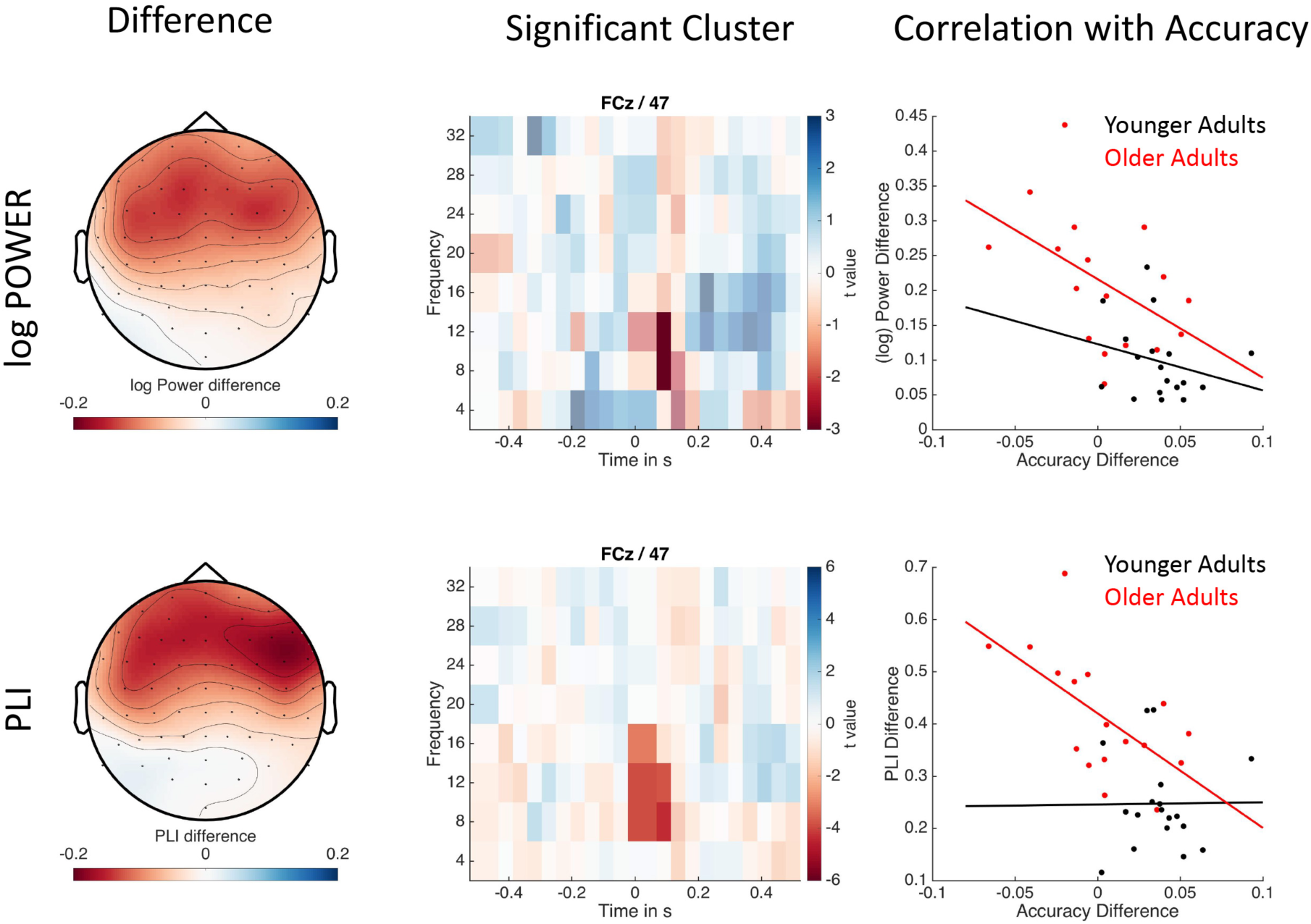
Age group differences in the cue effect as reflected in (log) power (upper panel) and PLI (lower panel) and their relation to the behavioral cue effect. Figure displays topography (left), significant cluster (highlighted) at FCz (middle) and the correlation with accuracy (right). Older adults show a stronger modulation of power and phase locking by the alerting cue that is related to a detrimental effect on accuracy.

Next, we investigated whether the higher power and phase locking to the cue in older age was related to age differences in the behavioral cue effect. The cue effect (thus, the cue – no-cue difference) on power, respectively PLI, and the cue effect on accuracy were indeed negatively related in the older adult sample (Power: *r* = −0.4769, *p* = 0.05, PLI: *r* = −0.611, *p* < 0.01). By contrast, there was no relation between the cue effects on accuracy and power or PLI in the younger adult sample (Power: *r* = −0.2553, *p* = 0.3065, PLI: *r* = 0.0097, *p* = 0.97). The correlation between PLI and accuracy within the older adult sample was significantly larger than the correlation in the young adult sample (one-sided test, *z* = −1.938, *p* = 0.026). Importantly, as can be seen in Fig. 5, the correlations between accuracy and both power and PLI in the older adults also indicate that despite overall age differences, those older adults that showed a cue-related performance benefit also showed PLI and power effects within the range of young adults.

## Discussion

The present study investigated age differences in power and inter-trial phase stability in response to an auditory warning cue in a phasic alerting paradigm. In both younger and older adults, the cue elicited similar levels of power increase over fronto-central areas in the alpha range. Differences between age groups were nevertheless found both in power and in phase locking. Particularly strikingly, older adults showed stronger phase locking than younger adults. In addition, higher phase locking was associated with a smaller behavioral alerting effect in the older age group. In fact, older adults with excessive phase locking to the alerting cue even experienced a detrimental effect in accuracy. These age differences in phase locking with simultaneous increases in power are indicative of a strong phase reset in older adults.

Higher phase locking in older than younger adults, similar to our results, has been reported previously in a visual working memory task (Sander et al., 2012) and an auditory oddball task (Müller et al., 2009). These age effects were suggested to indicate that sensory processing in older age is more automatic, stimulus-driven, and less easily modulated in a top-down, task-driven manner. As a consequence, older individuals’ system is less flexible and more responsive to external stimulation (Lindenberger and Mayr, 2014). The results of the present study are in line with this interpretation. Furthermore, the inverse relation between age and alerting effects on performance and phase locking implies that those older adults who showed a stronger stimulus-driven response to the cue benefitted less from it or were even impaired by it. Presumably, if the cue induced a strong resetting of the system, it hindered older adults from effectively using the warning signal to facilitate subsequent stimulus processing, as they could not disengage from processing the cue (or did so too slowly). In fact, the alerting cue may have served as distracter for a subsample of the older adults. The present findings lend indirect support to the notion that the duration and predictability of the time interval between the cue and subsequent stimulus is key to observing age differences (or not) (Zhou et al., 2011). Specifically, if disengagement from the cue is impaired in older adults, older adults are more likely to benefit from an alerting cue if time intervals between the cue and the stimulus are either consistent or otherwise long enough to reset the system to process the task stimulus (e.g., Haupt et al., 2018).

Notably, our results are in sharp contrast with a recent study that examined cue-related power and phase-locking effects in a visual working memory task in younger and older adults (Tran et al., 2016). This earlier study reported less consistent phase locking in older than in younger adults, which in turn predicted later visual memory performance. A number of important differences between this study and ours could explain the different results. First, Tran and colleagues (2016) presented alerting cues on each trial, whereas we presented them in only 50% of the trials. This design allowed us to quantify and compare both behavioral and neuronal alerting effects, that is, the difference in responses between cue and no-cue trials. In addition, while the cue-target interval varied randomly from trial to trial in our task, it was constant in the study by Tran and colleagues. Thus, besides bottom-up, reflexive alerting effects, voluntary temporal orienting (Weinbach and Henik, 2012), and potentially age differences therein (Chauvin et al., 2018; Zanto et al., 2011), may have contributed to their effects. If the target onset is temporally predictable, individuals are able to shift the phase of their alpha-band oscillations before target onset to optimize stimulus discrimination (Samaha et al., 2015). Accordingly, in Tran and colleagues’ (2016) study, young participants may have reset their phase to optimize stimulus processing in response to the cue. As the cue in our design was not informative of the exact time of stimulus occurrence, only phase resetting to the stimulus, but not to the cue onset would have been beneficial. In this case, strong phase locking to the cue may rather have hindered the older adults from imminently resetting their alpha oscillations to a phase that would be optimal for target stimulus processing shortly after the cue. Most importantly, the alerting cues used by the two studies differed. Whereas Tran and colleagues (2016) used transient (50 ms) color changes of the fixation cross, we used a loud tone as a cue. The latter is presumably more alerting and can hardly be missed due to eye blinks or attentional lapses, for example (both of which are more frequent in older than in young adults; see Carrière et al., 2010; McDowd and Shaw, 2000). Furthermore, the auditory alerting cue in the visual task allowed us to separate alerting effects from modality-specific sensory processing components that may overlap in time when both the cue and task stimulus are presented visually.

Finally, our results provide further evidence for the notion that the maintenance of a youth-like brain is key for healthy cognitive aging (Nyberg et al., 2012). Thus, despite an overall stronger phase locking in the older than in the younger adults, some of the older adults’ cue-related power and phase-locking effects were within the upper range of younger adults. Interestingly, such older adults with a more youth-like pattern also showed reliable performance benefits after the presentation of an alerting cue. Our finding is thus in line with the idea that high individual variability in older adults as reflected in the temporal dynamics of neural oscillations may be a valid marker for the functionality of attentional mechanisms (Mok et al., 2016). The functionality of attentional mechanisms, and more specifically, phasic alerting may in turn critically depend on brainstem structures (e.g., the noradrenergic and cholinergic system) that undergo strong changes over the lifespan (Robertson, 2013, 2014). Only recently, Dahl and colleagues (2018) provided evidence that older adults’ memory performance was positively related to a more youth-like integrity of the locus coeruleus. It is an intriguing possibility that individual differences in arousal may also be driven the integrity of the locus coeruleus. Thus, a promising road for future research is to understand how structural integrity of brainstem structures relate to individual differences in neural oscillations and alerting performance.

## Acknowledgments

The data for this study was collected within the *‘Center for Visual Cognition’* at the University of Copenhagen. Iris Wiegand was supported via a DFF & EU MSC-COFUND Mobilex Mobility Grant (1321-00039B) and an EU MSC Global Fellowship (702483). Myriam C. Sander was supported via Minerva Research Groups awarded by the Max Planck Society and by a grant from the German Research Foundation (DFG GZ BR 4918/2-1). We thank Julia Delius for editorial assistance.

## Footnotes

1 In no-cue trials, EEG triggers were set prior to stimulus onset using time intervals identical to the CTIs in cue trials (randomly drawn from a uniform distribution with a range of 240–330 ms using time steps of 10 ms).

## References

Bell, A. J., Sejnowski, T. J., 1995. An information-maximization approach to blind separation and blind deconvolution. Neural Comp., 7(6), 1129–1159. https://doi.org/10.1162/neco.1995.7.6.1129

Brown, S. B., Tona, K. D., van Noorden, M. S., Giltay, E. J., van der Wee, N. J., Nieuwenhuis, S., 2015. Noradrenergic and cholinergic effects on speed and sensitivity measures of phasic alerting. Behav. Neurosci., 129(1), 42–49. http://dx.doi.org/10.1037/bne0000030

Bundesen, C., 1990. A theory of visual attention. Psychol. Rev., 97(4), 523–547.

Carrière, J. S., Cheyne, J. A., Solman, G. J., Smilek, D., 2010. Age trends for failures of sustained attention. Psychol. Aging, 25(3), 569–574.

Chauvin, J. J., Gillebert, C. R., Rohenkohl, G., Humphreys, G. W., Nobre, A. C., 2016. Temporal orienting of attention can be preserved in normal aging. Psychol. Aging, 31(5), 442–455. http://dx.doi.org/10.1037/pag0000105

Coull, J. T., Nobre, A. C., Frith, C. D., 2001. The noradrenergic α2 agonist clonidine modulates behavioural and neuroanatomical correlates of human attentional orienting and alerting. Cereb. Cortex, 11(1), 73–84. https://doi.org/10.1093/cercor/11.1.73

Dahl, M. J., Mather, M., Duezel, S., Bodammer, N. C., Lindenberger, U., Kühn, S., Werkle-Bergner, M., 2018. Locus coeruleus integrity preserves memory performance across the adult life span. bioRxiv, 332098. https://doi.org/10.1101/332098

Deiber, M. P., Ibañez, V., Missonnier, P., Rodriguez, C., Giannakopoulos, P., 2013. Age-associated modulations of cerebral oscillatory patterns related to attention control. NeuroImage, 82, 531–546. https://doi.org/10.1016/j.neuroimage.2013.06.037

Delorme, A., Makeig, S., 2004. EEGLAB: An open source toolbox for analysis of single-trial EEG dynamics including independent component analysis. J. Neurosci. Meth., 134(1), 9–21. https://doi.org/10.1016/j.jneumeth.2003.10.009

Duncan, J., Bundesen, C., Olson, A., Humphreys, G., Chavda, S., Shibuya, H., 1999. Systematic analysis of deficits in visual attention. J. Exp. Psychol. Gen., 128(4), 450–478.

Festa-Martino, E., Ott, B. R., Heindel, W. C., 2004. Interactions between phasic alerting and spatial orienting: Effects of normal aging and Alzheimer’s disease. Neuropsychology, 18(2), 258–268. https://dx.doi.org/10.1037/0894-4105.18.2.258

Fernandez-Duque, D., Black, S. E., 2006. Attentional networks in normal aging and Alzheimer’s disease. Neuropsychology, 20(2), 133–143. https://dx.doi.org/10.1037/0894-4105.20.2.133

Folstein, M. F., Folstein, S. E., McHugh, P. R., 1975. Mini-mental state. A practical method for grading the state of patients for the clinician. J. Psych. Res., 12, 189–198.

Gamboz, N., Zamarian, S., Cavallero, C., 2010. Age-related differences in the attention network test (ANT). Exp. Aging Res., 36(3), 287–305. https://doi.org/10.1080/0361073X.2010.484729

Garrett, D. D., Samanez-Larkin, G. R., MacDonald, S. W., Lindenberger, U., McIntosh, A. R., Grady, C. L., 2013. Moment-to-moment brain signal variability: A next frontier in human brain mapping? Neurosci. Biobehav. Rev., 37(4), 610–624. https://doi.org/10.1016/j.neubiorev.2013.02.015

Hackley, S. A., Valle-Inclán, F., 2003. Which stages of processing are speeded by a warning signal?. Biol. Psychol., 64(1–2), 27–45. https://doi.org/10.1016/S0301-0511(03)00101-

Hanslmayr, S., Klimesch, W., Sauseng, P., Gruber, W., Doppelmayr, M., Freunberger, R., Pecherstorfer, T., 2005. Visual discrimination performance is related to decreased alpha amplitude but increased phase locking. Neurosci. Letters, 375(1), 64–68. https://doi.org/10.1016/j.neulet.2004.10.092

Hanslmayr, S., Klimesch, W., Sauseng, P., Gruber, W., Doppelmayr, M., Freunberger, R., Percherstorfer, T., Birbaumer, N., 2006. Alpha phase reset contributes to the generation of ERPs. Cereb. Cortex, 17(1), 1–8. https://doi.org/10.1093/cercor/bhj129

Hanslmayr, S., Aslan, A., Staudigl, T., Klimesch, W., Herrmann, C. S., Bäuml, K. H., 2007. Prestimulus oscillations predict visual perception performance between and within subjects. NeuroImage, 37(4), 1465–1473. https://doi.org/10.1016/j.neuroimage.2007.07.011

Haupt, M., Sorg, C., Napiórkowski, N., Finke, K., 2018. Phasic alertness cues modulate visual processing speed in healthy aging. Neurobiol. Aging, 70, 30–39. https://doi.org/10.1016/j.neurobiolaging.2018.05.034

Ishigami, Y., Eskes, G. A., Tyndall, A. V., Longman, R. S., Drogos, L. L., Poulin, M. J., 2016. The Attention Network Test-Interaction (ANT-I): reliability and validity in healthy older adults. Exp. Brain Res., 234(3), 815–827. https://doi.org/10.1007/s00221-015-4493-4

Jennings, J. M., Dagenbach, D., Engle, C. M., Funke, L. J., 2007. Age-related changes and the attention network task: An examination of alerting, orienting, and executive function. Aging Neuropsychol. Cogn., 14(4), 353–369. https://doi.org/10.1080/13825580600788837

Jensen, O., Mazaheri, A., 2010. Shaping functional architecture by oscillatory alpha activity: Gating by inhibition. Front. Hum. Neurosci., 4, 186. https://doi.org/10.3389/fnhum.2010.00186

Jung, T. P., Makeig, S., Westerfield, M., Townsend, J., Courchesne, E., Sejnowski, T. J., 2000. Removal of eye activity artifacts from visual event-related potentials in normal and clinical subjects. Clin. Neurophysiol., 111(10), 1745–1758. https://doi.org/10.1016/S1388-2457(00)00386-2

Klimesch, W., 1999. EEG alpha and theta oscillations reflect cognitive and memory performance: a review and analysis. Brain Res. Rev., 29(2–3), 169–195. https://doi.org/10.1016/S0165-0173(98)00056-3

Klimesch, W., Schack, B., Schabus, M., Doppelmayr, M., Gruber, W., Sauseng, P., 2004. Phase-locked alpha and theta oscillations generate the P1–N1 complex and are related to memory performance. Cogn. Brain Res., 19(3), 302–316. https://doi.org/10.1016/j.cogbrainres.2003.11.016

Klimesch, W., Sauseng, P., Hanslmayr, S., 2007. EEG alpha oscillations: The inhibition–timing hypothesis. Brain Res. Rev., 53(1), 63–88. https://doi.org/10.1016/j.brainresrev.2006.06.003

Kusnir, F., Chica, A. B., Mitsumasu, M. A., Bartolomeo, P., 2011. Phasic auditory alerting improves visual conscious perception. Conscious. Cogn., 20(4), 1201–1210. https://doi.org/10.1016/j.concog.2011.01.012

Lachaux, J. P., Rodriguez, E., Martinerie, J., Varela, F. J., 1999. Measuring phase synchrony in brain signals. Hum. Brain Map., 8(4), 194–208.

Lindenberger, U., Mayr, U., 2014. Cognitive aging: Is there a dark side to environmental support?. Trends Cogn. Sci, 18(1), 7–15. https://doi.org/10.1016/j.tics.2013.10.006

Maris, E., & Oostenveld, R., 2007. Nonparametric statistical testing of EEG and MEG-data. J. Neurosci. Meth., 164(1), 177–190. https://doi.org/10.1016/j.jneumeth.2007.03.024

Matthias, E., Bublak, P., Müller, H. J., Schneider, W. X., Krummenacher, J., Finke, K., 2010. The influence of alertness on spatial and nonspatial components of visual attention. J. Exp. Psychol. Hum. Percept. Perform., 36(1), 38–56. https://doi:10.1037/a0017602

McDowd, J. M., Shaw, R. J., 2000. Attention and aging: A functional perspective. In F. I. M. Craik and T. A. Salthouse (Eds.), The handbook of aging and cognition (pp. 221–292). Mahwah, NJ: Erlbaum.

Mok, R. M., Myers, N. E., Wallis, G., Nobre, A. C., 2016. Behavioral and neural markers of flexible attention over working memory in aging. Cereb. Cortex, 26(4), 1831–1842. https://doi.org/10.1093/cercor/bhw011

Müller, V., Gruber, W., Klimesch, W., Lindenberger, U., 2009. Lifespan differences in cortical dynamics of auditory perception. Develop. Sci., 12(6), 839–853. https://doi.org/10.1111/j.1467-7687.2009.00834.x

Nyberg, L., Lövdén, M., Riklund, K., Lindenberger, U., Bäckman, L., 2012. Memory aging and brain maintenance. Trends Cogn. Sci., 16(5), 292–305. https://doi.org/10.1016/j.tics.2012.04.005

Oostenveld, R., Fries, P., Maris, E., Schoffelen, J. M., 2011. FieldTrip: open source software for advanced analysis of MEG, EEG, and invasive electrophysiological data. Comput. Intel. Neurosci., 156869. https://doi:10.1155/2011/156869

Posner, M. I., Petersen, S. E., 1990. The attention system of the human brain. Annual review of neuroscience, 13(1), 25–42. http://dx.doi.org/10.1146/annurev.ne.13.030190.000325

Rabbitt, P. (1984). How old people prepare themselves for events which they expect. Attent. Perform., 10, 515–527.

Robertson, I. H., 2013. A noradrenergic theory of cognitive reserve: Implications for Alzheimer’s disease. Neurobiol. Aging, 34(1), 298–308. https://doi.org/10.1016/j.neurobiolaging.2012.05.019

Robertson, I. H., 2014. A right hemisphere role in cognitive reserve. Neurobiol. Aging, 35(6), 1375–1385. https://doi.org/10.1016/j.neurobiolaging.2013.11.028

Sander, M. C., Werkle-Bergner, M., Lindenberger, U., 2012. Amplitude modulations and inter-trial phase stability of alpha-oscillations differentially reflect working memory constraints across the lifespan. NeuroImage, 59(1), 646–654. https://doi.org/10.1016/j.neuroimage.2011.06.092

Sauseng, P., Klimesch, W. (2008). What does phase information of oscillatory brain activity tell us about cognitive processes?. Neurosci. Biobehav, Rev., 32(5), 1001–1013. https://doi.org/10.1016/j.neubiorev.2008.03.014

Sturm, W., Willmes, K., 2001. On the functional neuroanatomy of intrinsic and phasic alertness. NeuroImage, 14(1), 76–84. https://doi.org/10.1006/nimg.2001.0839

Sturm, W., De Simone, A., Krause, B. J., Specht, K., Hesselmann, V., Radermacher, I., Herzog, H., Tellmann, L., Müller-Gärtner, H.W., Willmes, K., 1999. Functional anatomy of intrinsic alertness: Evidence for a fronto-parietal-thalamic-brainstem network in the right hemisphere. Neuropsychologia, 37(7), 797–805. https://doi.org/10.1016/S0028-3932(98)00141-9

Tallon-Baudry, C., Bertrand, O., Delpuech, C., Pernier, J., 1996. Stimulus specificity of phase-locked and non-phase-locked 40 Hz visual responses in human. J. Neurosci., 16(13), 4240–4249. https://doi.org/10.1523/JNEUROSCI.16-13-04240.1996

Thiel, C. M., Fink, G. R., 2007. Visual and auditory alertness: modality-specific and supramodal neural mechanisms and their modulation by nicotine. J. Neurophysiol., 97(4), 2758–2768. https://doi.org/10.1152/jn.00017.2007

Thut, G., Nietzel, A., Brandt, S. A., Pascual-Leone, A., 2006. α-Band electroencephalographic activity over occipital cortex indexes visuospatial attention bias and predicts visual target detection. J. Neurosci., 26(37), 9494–9502. https://doi.org/10.1523/JNEUROSCI.0875-06.2006

Tran, T. T., Hoffner, N. C., LaHue, S. C., Tseng, L., Voytek, B., 2016. Alpha phase dynamics predict age-related visual working memory decline. NeuroImage, 143, 196–203. https://doi.org/10.1016/j.neuroimage.2016.08.052

Voytek, B., Kramer, M. A., Case, J., Lepage, K. Q., Tempesta, Z. R., Knight, R. T., Gazzaley, A., 2015. Age-related changes in 1/f neural electrophysiological noise. J. Neurosci., 35(38), 13257–12265. https://doi.org/10.1523/JNEUROSCI.2332-14.2015

Weinbach, N., Henik, A., 2012. Temporal orienting and alerting–the same or different?. Front. Psychol., 3, 236. https://doi.org/10.3389/fpsyg.2012.00236

Werkle-Bergner, M., Freunberger, R., Sander, M. C., Lindenberger, U., Klimesch, W., 2012. Inter-individual performance differences in younger and older adults differentially relate to amplitude modulations and phase stability of oscillations controlling working memory contents. NeuroImage, 60(1), 71–82. https://doi.org/10.1016/j.neuroimage.2011.11.071

Wiegand, I., Petersen, A., Finke, K., Bundesen, C., Lansner, J., Habekost, T., 2017a. Behavioral and brain measures of phasic alerting effects on visual attention. Front. Hum. Neurosci., 11, 176. https://doi.org/10.3389/fnhum.2017.00176

Wiegand, I., Petersen, A., Bundesen, C., Habekost, T., 2017b. Phasic alerting increases visual attention capacity in younger but not in older individuals. Vis. Cogn., 25(1–3), 343–357. https://doi.org/10.1080/13506285.2017.1330791

Zanto, T. P., Pan, P., Liu, H., Bollinger, J., Nobre, A. C., Gazzaley, A., 2011. Age-related changes in orienting attention in time. J. Neurosci., 31(35), 12461–12470. https://doi.org/10.1523/JNEUROSCI.1149-11.2011

Zhou, S. S., Fan, J., Lee, T. M., Wang, C. Q., Wang, K., 2011. Age-related differences in attentional networks of alerting and executive control in young, middle-aged, and older Chinese adults. Brain Cogn., 75(2), 205–210. https://doi.org/10.1016/j.bandc.2010.12.003

